# Potential survival strategies of novel comammox and nitrite-oxidizing *Nitrospira* present in a reactor treating high-ammonia brackish landfill leachate

**DOI:** 10.1101/2023.12.27.573385

**Authors:** Shohei Yasuda, Alejandro Palomo, Barth F. Smets, Akihiko Terada

## Abstract

Nitrification is mediated by numerous different microorganisms, but knowledge of their ecophysiologies is insufficient. Leachate in the late stages of landfill operation provides a brackish environment with a high ammonia concentration, and methanol is added as an electron donor for denitrification. Such a unique environment may contain novel nitrifiers. Here, we present metagenomic analysis of the microbiome from a closed landfill leachate treatment facility to investigate the identity and functions of nitrifiers. Using a genome-centric approach with metagenomic analysis, we retrieved draft genomes for a novel complete ammonia-oxidizing (comammox) bacterium *Nitrospira* LAS72; and canonical *Nitrospira* LAS18, clustered within a novel sub-lineage VII of *Nitrospira*; *Candidatus* Nitrosocosmicus LAS21 and *Nitrosarchaeum* LAS73, belonging to the ammonia-oxidizing archaea (AOA). This is the first evidence of comammox *Nitrospira* in a high-ammonia-containing brackish environment. Canonical ammonia-oxidizing bacteria were not detected. Given the brackish environment and supplementation of methanol used in the facility, we also investigated the methanol metabolism of these nitrifiers and their potential to produce compatible solutes as osmoprotectants. Uniquely among *Nitrospira*, comammox *Nitrospira* LAS72 possesses genes associated with formaldehyde reductase and glycine betaine biosynthesis. Thus, *Nitrospira* LAS72 may proliferate because of the availability of formaldehyde upstream of carbon metabolism and adapt to fluctuating osmotic pressure by producing a variety of compatible solutes. The discovery of this novel comammox *Nitrospira,* and canonical *Nitrospira* forming a new sub-lineage VII in an ammonia-concentrated brackish environment broadens our knowledge of the diversity and functions of nitrifying microorganisms.

## Introduction

Nitrification proceeds through various microbial processes depending on the environment. Ammonia-oxidizing archaea (AOA), ammonia-oxidizing bacteria (AOB), and nitrite-oxidizing bacteria (NOB) have been known for a long time as microorganisms involved in nitrification. Identities and functions of these nitrifiers in activated sludge [1, 2], lakes [3], rivers [4], and estuaries that are saline environments [5, 6] have been investigated. Generally, AOB and AOA, also known as ammonia-oxidizing microorganisms (AOM), drive ammonia oxidation and convert ammonia to nitrite, which is then further oxidized to nitrate by NOB, including the genus *Nitrospira* [7]. In recent years, however, the existence of complete ammonia-oxidizing (comammox) *Nitrospira* has been reported; these bacteria undertake complete conversion of ammonia to nitrate via nitrite [7, 8]. *N. inopinata* was first isolated as a comammox bacterium from an oligotrophic environment [9].

The genus *Nitrospira* is diverse, with seven types of lineages; comammox *Nitrospira* belong to lineage II. Their ecological niche is differentiated by dissolved oxygen and nitrite concentrations [10]. Comammox *Nitrospira* within lineage II are further grouped into clades A and B by phylogenetic classification of the ammonia monooxygenase [7, 11]. Combining amplicon sequence analysis using the ammonia monooxygenase (*amoA*) gene derived from comammox *Nitrospira* and metagenomic analysis, comammox *Nitrospira* have been detected in many environments, including rivers, forest soils, paddy soils, rice rhizosphere, drinking water treatment plants, rapid sand filters, wastewater treatment plants (WWTPs), and particulate matter (PM_2.5_) in the atmosphere, suggesting their presence in a wide range of oligotrophic environments (0.001–65 mg-NH_4_^+^-N/L) [12–19]. Comammox detection in brackish water areas [20] and sediments of mangrove forests close to the salt concentration of seawater (0.5%–3%) has been reported by detection of the *amoA* gene [21]. Meanwhile, our previous work employed gene-centric metagenomic analysis of the microbiome inhabiting a landfill leachate treatment plant, unveiling the presence of comammox *Nitrospira* in a brackish (salinity 1.23%) and high-ammonia (117.80 ± 30.13 mg-NH_4_^+^-N/L) environment [22]. However, these analyses do not comprehensively describe the functions of comammox *Nitrospira*. In particular, the reasons for their presence in a high-salinity and -ammonia environment, their relative abundance, and their contribution to ammonia oxidation remain unknown. Therefore, here, we aimed to deepen our understanding of the physiological ecology of comammox *Nitrospira* in the unique environment of the microbiome of a leachate treatment plant.

Landfill leachate has unique and complex compositions and constituents, generated by the infiltration of rainwater into landfill, which depend on the waste composition, landfill age, climatic conditions, and the hydrogeology of the landfill [23]. The organic matter reclaimed in landfill is anaerobically degraded and converted into methane and ammonia by microorganisms. In the later stage of the landfill (after several decades), leachate includes not only low levels of organic matter and a high concentration of ammonia, but also high salinity derived from incineration residues [24, 25]. Leachate treatment plants install a system to perform nitrification/denitrification, and methanol is often added as an electron donor to supplement the low-concentration organic matter to promote denitrification [22]. Biomass in the treatment plant was recycled from a sedimentation tank to a reaction tank. It has been subjected to feast and famine conditions because of achieving high efficiencies in organic carbon and nitrogen removal [22]. From a different perspective, the treatment plant can be likened to conducting large-scale enrichment culture on an actual scale, and it has been in operation for several decades since the commencement of landfill leachate treatment. The long-term enrichment culture selected microorganisms that could gradually adapt to this unique environment. However, the 16S rRNA gene amplicon sequencing and gene-centric metagenome analysis, applied in previous studies [22, 26] to investigate microbial community compositions and functions, have not fully succeeded in identifying nitrifiers, determining their abundance, and characterizing their individual genomic features.

This study used genome-centric metagenomics to investigate nitrifiers in the microbiome treating landfill leachate in a brackish environment with high ammonia concentration and added methanol as a carbon source. To this end, we asked the following research questions:

- What are the dominant nitrifiers in this unique environment?
- What are their unique genomic features?
- Is it possible to obtain a draft comammox *Nitrospira* genome, and, if so, what type is it and what genetic characteristics does it have?

Our previous gene-centric analysis of the same microbial community revealed the presence of *Nitrospira* and *Thaumarchaeota*, indicating the potential for acquisition of draft genomes for *Nitrospira*, including comammox strains, and AOA [22]. Hybrid sequencing with long and short reads, and integration of three binning algorithms to calculate an optimized set of bins from a single assembly, were performed to acquire high-quality metagenomic assembled genomes (MAGs) of microorganisms involved in nitrification. By understanding the genomic features of comammox *Nitrospira* in an unexplored eutrophic brackish environment with relatively high ammonia concentration, this study discovered novel comammox *Nitrospira* phylogeny and functionality, broadening our knowledge of nitrification mechanisms in high-ammonia and saline environments.

## Materials and Methods

### Sampling, DNA extraction, and purification

Biomass collected from a leachate treatment plant in July 2018 (Tokyo, Japan) was used in this study; it was the same material as used in a previous study [22]. Briefly, this biomass was taken from a sequencing batch reactor (SBR) intermittently receiving a mixture of landfill leachate and groundwater. The treated water was intermittently transferred to an aeration tank for stabilization by a pump, followed by a sedimentation tank to separate biomass and treated water. The sedimented biomass was returned to the SBR to retain biomass concentration. The detailed system configuration is shown elsewhere [22]. The leachate (average flow rate: 74.2 ± 40.0 m^3^/day) containing NH_4_^+^ at 151.31 ± 49.53 mg-N/L and chemical oxygen demand (COD) at 43.9 ± 9.2 mg-COD/L was mixed with groundwater (13.2 ± 11.1 m^3^/day) in a flow equalization tank, located before the SBR. The tank also received methanol as an external carbon source, resulting in an average NH_4_^+^ concentration of 117.80 ± 30.13 mg-N/L and an average COD of 229.47 ± 93.1 mg-COD/L. The treated water contained average COD and NH_4_^+^ concentrations below 10.0 mg-COD/L and 0.05 mg-N/L, respectively. Twenty liters of the biomass taken during an aeration period of the intermittent aeration tank was immediately transferred to the laboratory, and 50 mL of the thoroughly mixed biomass was stored at −20°C until use. DNA extraction was performed by the phenol–chloroform method [27], and contaminating RNA was degraded by using RNase A (TaKaRa Bio, Inc., Shiga, Japan).

### Hybrid assembly

Hybrid metagenomic assembly with short and long reads was performed to attain high-quality MAGs. For the long-read sequencing procedure, the library was prepared with a 1D ligation sequencing kit (SQK-LSK-109; Oxford Nanopore Technologies PLC, Oxford, UK) without a fragmentation procedure and sequenced on the MinION Mk1B instrument using a flow cell (FLO-MIN106; Oxford Nanopore Technologies). The sequence data were base called with Guppy ver. 3.6.1 (Oxford Nanopore Technologies), and the sequence quality was confirmed with NanoPlot ver. 1.20.0 [28]. The adaptor sequence was removed using Porechop ver. 0.2.4 (https://github.com/rrwick/Porechop), and the low-quality reads (Q6), header (75 bp), and short reads (<=1,000 bp) were trimmed with NanoFilt ver. 2.2.0 [28]. In short-read sequencing, two indices were used and sequenced twice to ensure barcode diversity. The libraries were prepared with the MGIEasy universal DNA library prep set according to the manufacturer’s protocol (MGI Tech, Shenzhen, China), and 150-bp paired-end sequencing was performed using DNBSEQ-G400FAST (MGI Tech). Adapter sequences and low-quality reads (Q30) were removed using Trim Galore ver. 0.6.5 (http://www.bioinformatics.babraham.ac.uk/projects/trim_galore/). The consensus sequence was assembled with Megahit ver. 1.2.9 [29]. The short reads and the trimmed long reads were mapped to the assembly, and the obtained connectivity between the contigs and read coverage information was used to accurately cluster contigs into genomes with OPERA-MS ver. 0.8.2 [30] using minimap2 ver. 2.17 [31] as the long read mapper. The hybrid sequence quality was confirmed with QUAST ver. 5.0.2 [32].

### Auto and manual binning

Anvio ver. 6.2 [33] was used as an analytical platform. A contigs database was created from the hybrid sequences. After an SAM file was made with BWA-MEM ver. 0.7.17 [34], conversion of the SAM file to a BAM file and sorting and indexing the BAM file were implemented in SAMtools ver. 1.9 [35]. Annotations with GhostKOALA [36], a comprehensive collection of protein domains and families (Pfam) [37], Clusters of Orthologous Groups [38], and single-copy core genes were conducted using the Anvio platform. The clustering of contigs was implemented with binning software with CONCOCT ver. 0.4.2 [39], Metabat2 ver. 2.13 [40], and maxbin2 ver. 2.2.5 [41], and the results were integrated with DAS_Tool ver. 1.1.2 [42]. Finally, manual binning was performed to attain highly refined bins. The completeness and contamination of each bin were confirmed with CheckM ver. 1.0.7 [43], and the draft genome qualities were assessed according to the genome reporting standards for MAGs [44].

### MAG coverage, and *amoA* and *hao* sequence analysis

CoverM ver. 0.6.1 (https://github.com/wwood/CoverM) was used for calculating MAG coverage and each relative abundance among total populations was obtained. To assess the accuracy of coverage, each gene sequence of *amoA* and hydroxylamine dehydrogenase (*hao*) sequence was confirmed using Vsearch ver. 2.17.1 [45] with the *amoA* and *hao* database for different species/genera/clades of AOA, AOB, and comammox *Nitrospira* obtained from UniProt (https://www.uniprot.org/).

### Genomic and functional annotation of MAGs

Genomic annotations of MAGs were performed with DFAST ver. 1.2.13 [46] with parameters minimum sequence length 200, first 100 bp offset, and aligned with blastp [47]. To understand the taxonomies for MAGs, 16S rRNA gene sequences were extracted and collated using the blastn database. Functional annotation was implemented via the Kyoto Encyclopedia of Genes and Genomes (KEGG) [48] using BlastKOALA ver. 2.2 [36] to understand functional traits for central metabolism, nitrification, and other metabolic pathways (including methanol metabolism, because methanol was the primary carbon source in the landfill leachate treatment facility). The salt concentration fluctuates in the SBR because of the mixing of leachate and groundwater and the dewatering after wastewater treatment, which requires the microbiome to quickly respond to changes in osmotic pressure. Therefore, functional genes related to osmoprotectant synthesis and related transporters were also investigated. Genes associated with urea transport and decomposition were investigated to explore the ability of the system to use heterotrophic bacteria- and detritus-derived urea in nitrogen-limited conditions after nitrification/denitrification.

### Phylogenic tree for *Nitrospira*

Thirty-eight *Nitrospira* genomes downloaded from NCBI were used to construct a phylogenic tree with GTDB-Tk ver. 2.2.1 [49] using the *de novo* workflow with a set of 120 translated universal single-copy genes and the genome taxonomy database (GTDB) [50]. A maximum-likelihood tree was built from concatenated alignments using RAxML ver. 8.2.11 [51] with 400 rapid bootstraps determined by autoMRE, and the LG likelihood model of amino acid substitution with gamma distributed rates (-m PROTGAMMALGF). The substitution models were selected using ProtTest ver. 3.4.2 [52]. The tree was rooted with two *Leptospirillum* species as an outgroup. The online web tool from the Interactive Tree of Life (iTol) ver. 3 [53] was used for visualization.

### Phylogenic affiliation and the average amino acid identity (AAI) comparison for a novel nitrite-oxidizing *Nitrospira*

Phylogenic tree with *Nitrospira* genomes downloaded from NCBI showed that canonical *Nitrospira* LAS18 might represent a novel lineage (**Fig. 1**). To deepen our knowledge for LAS18, further phylogenic affiliation with all *Nitrospira* genomes (downloaded in November 2023) using above mentioned methods, average amino acid identity (AAI) comparison, and 16S rRNA gene amplicon sequencing analysis were processed. The AAI between pair of genomes was calculated using EzAAI ver. 1.2.2 [54]. 16S rRNA gene sequences of LAS18 incorporating the same from 168 *Nitrospira* species downloaded from NCBI were aligned using MUSCLE (Edgar, 2004). The phylogenetic tree was inferred by using the Maximum Likelihood method and Tamura-Nei model in MEGA X [55] with a bootstrap test of 100 replications.

**Fig. 1.**
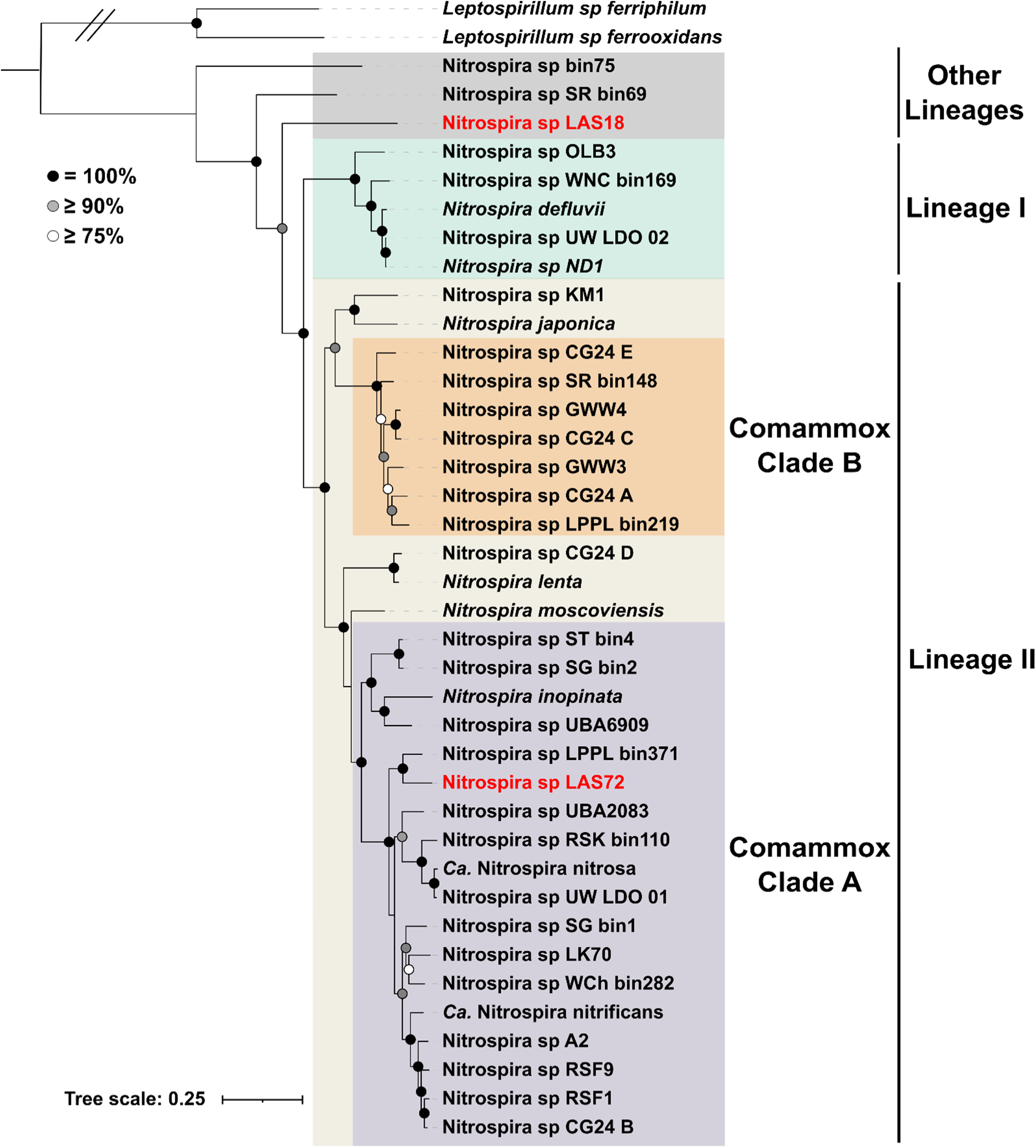
Genome based taxonomic classification for comammox *Nitrospira* LAS72 and canonical *Nitrospira* LAS18 affiliated with 38 *Nitrospira* genomes downloaded from the NCBI database. Comammox *Nitrospira* LAS72 and canonical *Nitrospira* LAS18 are highlighted in red.

### Pangenome and average nucleotide identity (ANI) analysis for *Nitrospira*

Pangenome analysis for the genus *Nitrospira* was implemented to identify core and singleton genes. Twenty high-quality *Nitrospira* genomes with completeness >90% and contamination <10% were retrieved from the NCBI database (downloaded in June 2021) (**Table S1**), and two *Nitrospira* MAGs obtained in this study, *i.e.*, for canonical *Nitrospira* LAS18 and comammox *Nitrospira* LAS72, were compared with them. The analysis was performed with the Anvio platform using minbit parameter 0.5 in ITEP ver. 1.1 [56], and inflation parameter 10.0 in the MCL algorithm [57] to identify clusters in amino acid sequence similarity search results. The Blastp (https://blast.ncbi.nlm.nih.gov/Blast.cgi) was used for calculating the similarity between all the gene calls in all the genomes. The ANI of each MAG was confirmed with pyANI ver. 0.2.10 [58].

#### KEGG-based singleton sequence analysis for canonical *Nitrospira* LAS18 and comammox Nitrospira LAS72

To understand unique metabolic pathways in canonical *Nitrospira* LAS18 and comammox *Nitrospira* LAS72, singleton genes that could be annotated with KEGG were investigated. *Nitrospira* genomes obtained from the NCBI database were annotated with GhostKOALA ver. 2.1 [36]. After comparing these genomes with the genomes of canonical *Nitrospira* LAS18 and comammox *Nitrospira* LAS72, genes unique to canonical *Nitrospira* LAS18 and comammox *Nitrospira* LAS72 were extracted. Genes different from the extracted ones but having the same function instrumental in the same metabolic pathway were investigated, and if any other *Nitrospira* possessed such a gene, the pathway was eliminated from the unique and independent pathways. For all analyses in this study, default parameters were used unless otherwise specified.

## Results

### Assembly and binning

Hybrid metagenomic assembly was conducted from short and long reads from the microbiome in the landfill leachate treatment plant (**Tables S2** and **S3**). In the short-read sequencing, 19.28 Gbp of total base pairs were obtained, and there were 64.26 M total reads. In the long-read sequencing, 25.6 Gbp and 3.2 M reads were obtained. These sequences were hybridized to give 807.6 Mbp, 468,765 contigs, and 43,606 bp for N50. Considering MAGs other than of nitrifiers, 97 high- and middle-level quality MAGs (completeness >50%, contamination <10%) were reconstructed. *Methyloceanibacter*, a methanol assimilating bacterium, was the most abundant in the whole microbial community.

### Microorganisms related to nitrification

Considering nitrifiers, MAGs for two types of AOA and two types of *Nitrospira* were obtained (**Table 1**). No canonical AOB MAGs or AOB *amoA* genes were detected. MAGs LAS18 and LAS72, belonging to *Nitrospira*, had >90% completeness and <5% contamination. They encoded 5S, 16S, and 23S rRNA sequences, and ≥18 tRNA sequences, indicating very high quality. LAS18 and LAS72 contain genes encoding nitrite oxidoreductase (*nxr*). The *amo* and *hao* operons are present in LAS72 but absent from LAS18, suggesting that LAS72 completely oxidizes ammonia to nitrate, while LAS18 oxidizes nitrite to nitrate. Phylogenetic analysis revealed that LAS72 is affiliated with *Nitrospira* linage II comammox clade A, while LAS18 does not belong to any other *Nitrospira* lineages, suggesting that this MAG represents a new lineage of the genus *Nitrospira* (**Fig. 1**). LAS21 and LAS73 belong to the AOA and were taxonomically classified as *Candidatus* Nitrosocosmicus (MAG completeness 81.58%, contamination 1.32%), and *Nitrosarchaeum* (completeness 67.11%, contamination 0%), respectively (**Table 1**).

**Table 1.** Characteristics and general features for MAGs information for canonical *Nitrospira* LAS18, comammox *Nitrospira* LAS72, *Candidatus* Nitrosocosmicus LAS21, and *Nitrosarchaeum* LAS73. The genome quality was judged with the genome reporting standards for MAGs proposed by Bowers et al, 2017. The coverage was the percentage of mapped reads to each MAG. The relative abundance was a percentage of each MAG in the total community.

|  | LAS18 | LAS72 | LAS21 | LAS73 |
| --- | --- | --- | --- | --- |
| Classification | NOB | Comammox | AOA | AOA |
| Taxonomy | *Nitrospira* | *Nitrospira* | *Candidatus Nitrosocosmicus* | *Nitrosarchaeum* |
| Completeness [%] | 95.91 | 93.18 | 83.98 | 70.39 |
| Contamination [%] | 3.69 | 3.38 | 1.94 | 0 |
| Genome quality | High | High | Medium | Medium |
| 5S rRNA gene | + | + | + | + |
| 16S rRNA gene | + | + | + | + |
| 23S rRNA gene | + | + | + | + |
| 18/20 tRNA | + | + | - | - |
| Total length [Mbp] | 4.35 | 3.84 | 2.14 | 1.33 |
| Num contigs | 11 | 44 | 31 | 5 |
| N50 [Mbp] | 4.20 | 0.12 | 0.11 | 0.36 |
| GC content [%] | 58.8 | 54.9 | 34.9 | 33.1 |
| Coverage | 82.6 | 13.2 | 63.4 | 13.0 |
| Relative Abundance [%] | 2.17 | 0.25 | 1.62 | 0.33 |

*Candidatus* Nitrosocosmicus LAS21 (The relative abundance in the total community was 1.62%) had the highest relative abundance among the AOM, and physiologically canonical *Nitrospira* LAS18 (2.17%) was the dominant nitrite-oxidizing microbe (**Table 1** and **Fig. 2**). To confirm these observations and rule out the presence of AOB, which were not revealed by the binning approach, in the community, we obtained *amoA* and *hao* sequences derived from AOA, AOB, and comammox *Nitrospira* from a public database, and recruited all reads mapping to the *amoA* sequences. The results were in line with those obtained based on the MAG abundances (**Table 1** and **Fig. 2**).

**Fig. 2.**
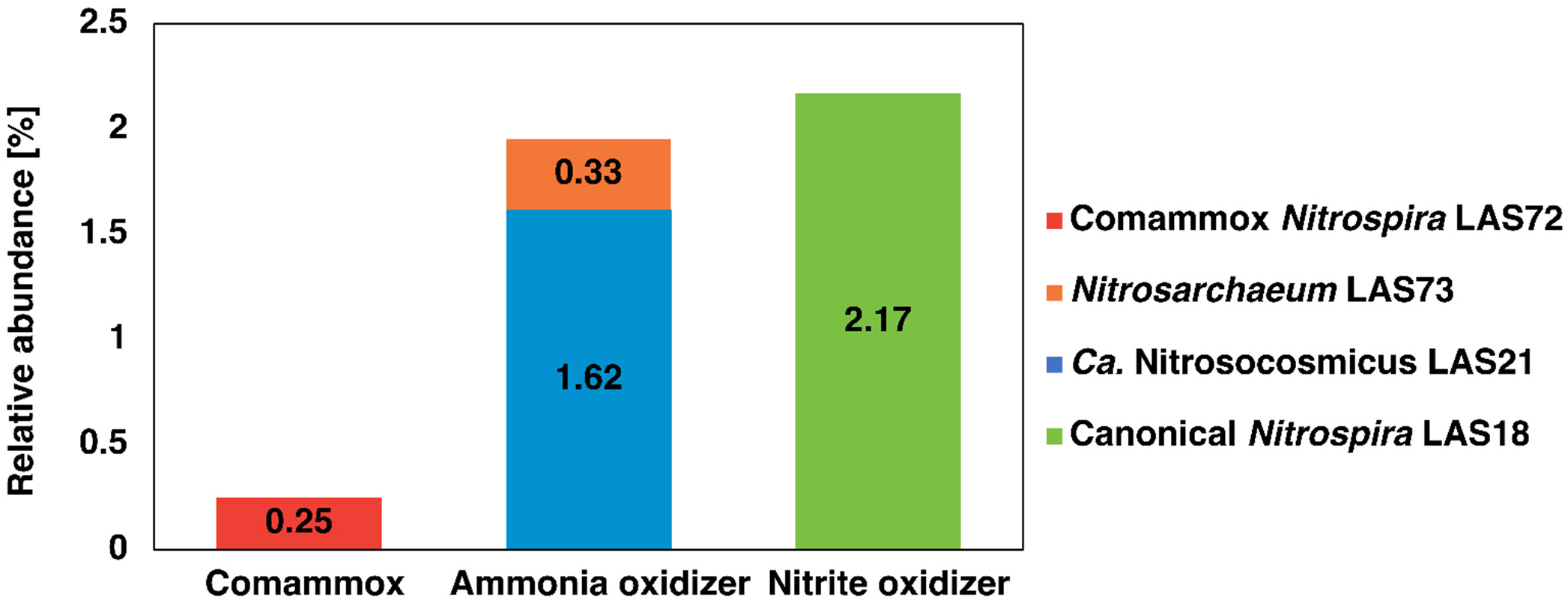
Relative abundances of four nitrifiers including canonical *Nitrospira* LAS18, comammox *Nitrospira* LAS72, *Candidatus* Nitrosocosmicus LAS21, and *Nitrosarchaeum* LAS73 in the total community.

### A novel nitrite-oxidizing *Nitrospira*

Phylogenic analysis indicated that the canonical *Nitrospira* LAS18 in the high ammonia brackish environment might represent a novel lineage (**Fig. 1**). To further support this finding, all *Nitrospira* genomes uploaded to the NCBI database with completeness above 70% and contamination below 5% were used for phylogenic affiliation and AAI comparison. Additionally, a phylogenetic tree of LAS18, incorporating 16S rRNA gene sequences from 168 *Nitrospira* species downloaded from NCBI, was constructed. Both phylogenetic trees, based on 120 marker proteins (**Fig. 3**) and 16S rRNA genes (**Fig. S2**), indicate that LAS18 stands apart from other lineages, possibly aligning with the new lineage VII, where *Nitrospira tepida* serves as the culture/genome representative [59]. The AAI value comparing LAS18 and *N. tepida* was determined to be 69.49% (**Fig. S1**), while the 16S rRNA gene sequence similarity between the two exhibited a value of 95.19% (**Fig. S2**). According to established criteria, AAI values of 65%–95% are indicative of different species, and for the 16S rRNA gene, values of 95%–98.6% typically denote the same genus [60]. In this context, LAS18 and *N. tepida* appear to belong to the same genus but different species. In addition, supported by the placement of these two genomes in the phylogenetic tree of the *Nitrospira* genus (**Fig. 3** and **Fig. S2**), LAS18 is likely part of the same lineage as *N. tepida* but forms a distinct sub-lineage within lineage VII. Notably, *Nitrospira* LAS18 is the only representative with a sequenced genome in that sub-lineage.

**Fig. 3.**
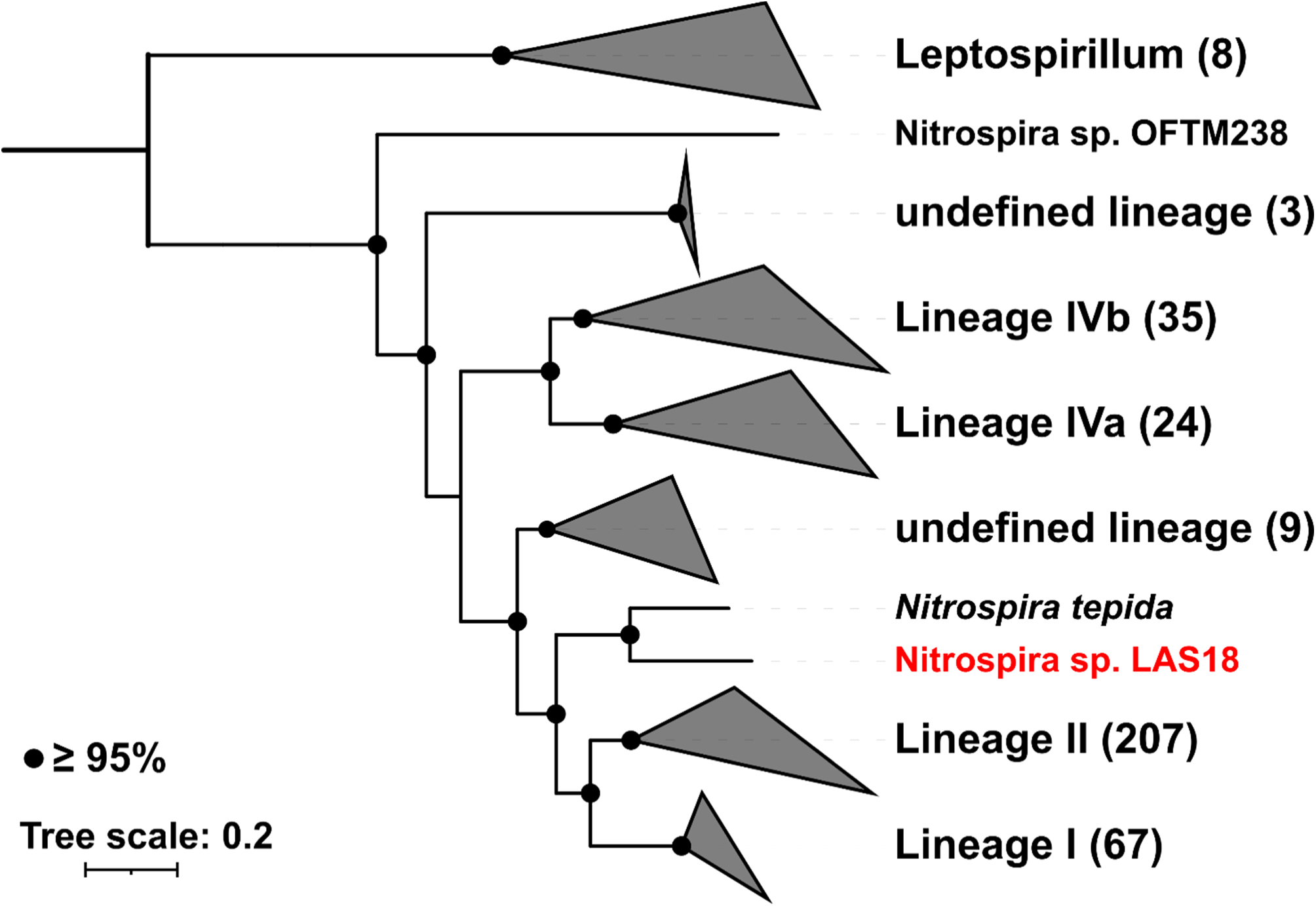
Genome based taxonomic classification for canonical *Nitrospira* LAS18 affiliated with 351 *Nitrospira* genomes downloaded from the NCBI database. Canonical *Nitrospira* LAS18 is highlighted in red. The number of genomes in each lineage was found inside of parenthesis.

### Functional analysis of *Nitrospira* LAS18, comammox *Nitrospira* LAS72, and AOA

By KEGG-mapping the amino acid sequences of canonical *Nitrospira* LAS18, comammox *Nitrospira* LAS72, and the AOA MAGs to reference metabolisms, the presence of functional genes and metabolic pathways was confirmed (**Table 2** and **S4**). Regarding ammonia oxidation, comammox *Nitrospira* LAS72 harbored *amoABC*, the genes for canonical subunits of ammonia monooxygenase, but these genes were not detected in the *Nitrospira* LAS18 and *Nitrosarchaeum* LAS73 MAGs. *Candidatus* Nitrosocosmicus LAS21 possessed *amoB*, the gene for a subunit of potentially active site of archaeal ammonia monooxygenase. The non-detection of *amoABC* in *Nitrosarchaeum* LAS73 is probably because of the incompleteness of the MAG (70.39% completeness). Regarding hydroxylamine oxidation, LAS72 possessed *hao*, encoding hydroxylamine oxidoreductase. Canonical *Nitrospira* LAS18 also possessed an analogous gene to *hao*, but probably encoding a cytochrome without *hao* function because the similarity with *hao* present in comammox *Nitrospira* and canonical AOB was only 30%. Considering nitrite oxidation, canonical *Nitrospira* LAS18 and comammox *Nitrospira* LAS72 harbored *nxrAB*. Canonical *Nitrospira* LAS18 and comammox *Nitrospira* LAS72 harbored *amt* and *slc42a-1*, encoding ammonium transporters. For urea degradation, canonical *Nitrospira* LAS18, comammox *Nitrospira* LAS72, and *Nitrosarchaeum* LAS73 had *ureABCDFG*. As urea transporters, canonical *Nitrospira* LAS18 possessed *urtABCDE* and comammox *Nitrospira* LAS72 had *urtCDE*. Regarding nitrite reduction, all four genomes contained *nirK*. Considering nitric oxide reduction, canonical *Nitrospira* LAS18 and comammox *Nitrospira* LAS72 harbored *norD* and *norQ*, encoding accessory proteins of nitric oxide reductase; however, they lack *norB* and *norC*, encoding NorBC, the catalytic subunits of nitric oxide reductase. *Candidatus* Nitrosocosmicus LAS21 possessed *norQ*. None of the four MAGs contained the *nosZ* gene associated with nitrous oxide reduction.

**Table 2.**
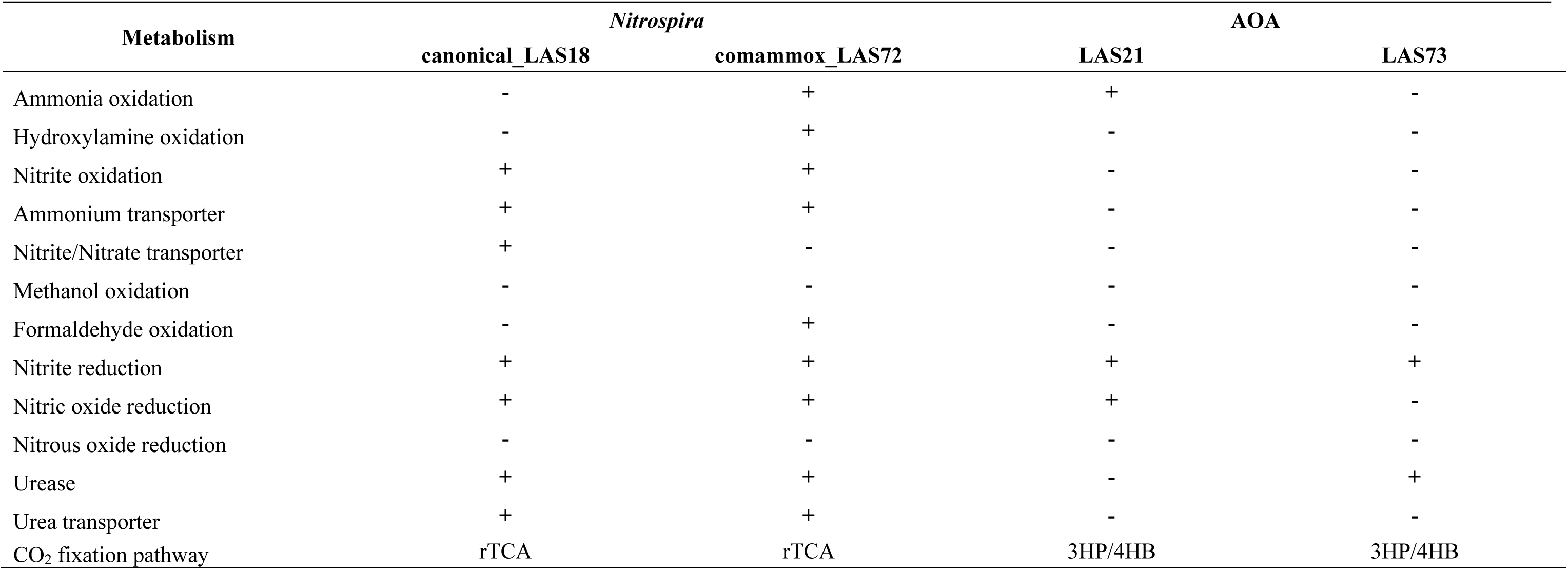
Presence or absence of metabolisms for four nitrifiers obtained in this study. For CO_2_ fixation, the assumed pathway cycle is shown. The abbreviations include rTCA: reverse TCA cycle, and 3HP/4HB: 3-hydroxypropionate/4-hydroxybutyrate cycle.

| Metabolism | *Nitrospira* |  | AOA |  |
| --- | --- | --- | --- | --- |
|  | canonical_LAS18 | comammox_LAS72 | LAS21 | LAS73 |
| Ammonia oxidation | - | + | + | - |
| Hydroxylamine oxidation | - | + | - | - |
| Nitrite oxidation | + | + | - | - |
| Ammonium transporter | + | + | - | - |
| Nitrite/Nitrate transporter | + | - | - | - |
| Methanol oxidation | - | - | - | - |
| Formaldehyde oxidation | - | + | - | - |
| Nitrite reduction | + | + | + | + |
| Nitric oxide reduction | + | + | + | - |
| Nitrous oxide reduction | - | - | - | - |
| Urease | + | + | - | + |
| Urea transporter | + | + | - | - |
| CO<sub>2</sub> fixation pathway | rTCA | rTCA | 3HP/4HB | 3HP/4HB |

Regarding the carbon fixation metabolism in the four nitrifying genomes, comammox *Nitrospira* LAS72 and canonical *Nitrospira* LAS18 contained most of the genes for the reductive tricarboxylic acid cycle (rTCA) cycle, except for the genes encoding aconitate hydratase, isocitrate dehydrogenase, and succinyl-CoA synthetase in LAS72, and the gene encoding oxoglutarate dehydrogenase in LAS18. In contrast, the AOA MAGs, LAS73 and LAS21, contained genes for the 3-hydroxypropionate/4-hydroxybutyrate carbon fixation pathway, as previously reported [61, 62]. None of the four nitrifiers carried the gene encoding methanol dehydrogenase (associated with methanol oxidation). However, comammox *Nitrospira* LAS72 possessed the *fdhA* gene encoding glutathione-independent formaldehyde dehydrogenase, suggesting that LAS72 assimilates formaldehyde.

### Pangenome analysis of *Nitrospira*

A comparative genomic analysis was performed to compare the genomic uniqueness of canonical *Nitrospira* LAS18 and comammox *Nitrospira* LAS72 with 20 high-quality *Nitrospira* genomes, including 12 comammox and 8 canonical *Nitrospira*, downloaded from NCBI (**Table S1**). On the basis of ANI analysis, both canonical *Nitrospira* LAS18 and comammox *Nitrospira* LAS72 had a similarity of <91% with respect to all other genomes, indicating that they constitute unique species (**Fig. 4** and **Fig. S3**). Pangenome analysis identified 22,744 gene clusters consisting of 84,824 genes, with 3.1% (700 gene clusters consisting of 16,392 genes) being core gene clusters and 59.1% (13,438 gene clusters consisting of 13,838 genes) being singleton gene clusters. The numbers of singleton genes in canonical *Nitrospira* LAS18 and comammox *Nitrospira* LAS72 were 1,508 and 675, respectively. To help understand the unique genetic characteristics of these bacteria, KEGG-annotated genes possessed only by canonical *Nitrospira* LAS18 or comammox *Nitrospira* LAS72 were analyzed (**Table S5**). Canonical *Nitrospira* LAS18 possessed 41 KEGG-annotated characteristic genes, including the *CLCN2* gene encoding chloride channel 2, the *GSK3A* gene encoding glycogen synthase kinase 3 alpha, *glpK* encoding glycerol kinase, and *ompR*, the phosphate regulon response regulator related to osmolarity. Comammox *Nitrospira* LAS72 harbored 18 KEGG-annotated singleton genes, including *gsmt-sdmt* encoding glycine/sarcosine N-methyltransferase and sarcosine/dimethylglycine N-methyltransferase associated with the production of glycine betaine from glycine, and *opuD*, which encodes a glycine betaine transporter.

**Fig. 4.**
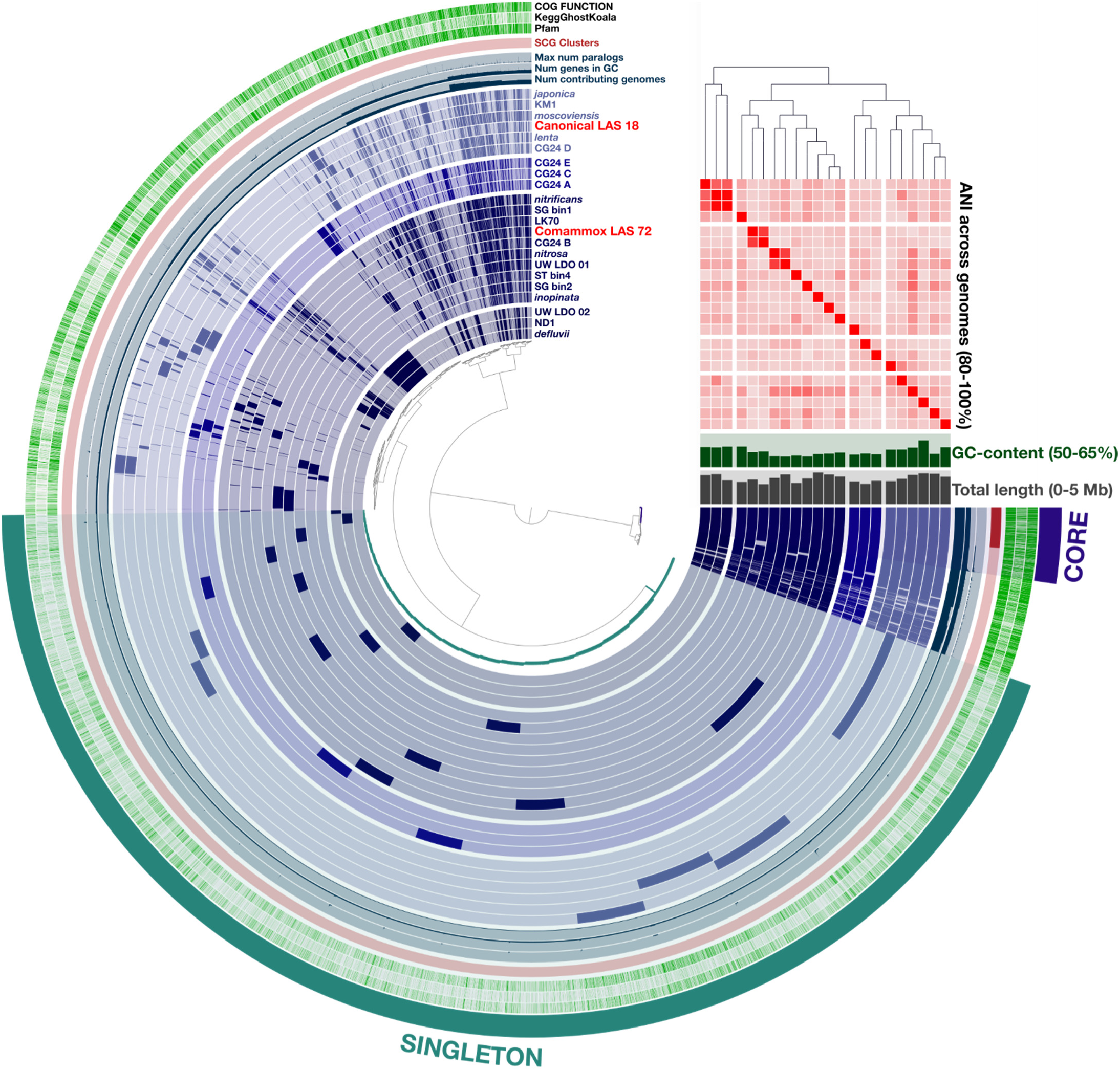
*Nitrospira* pangenome using 22 *Nitrospira* genomes including canonical *Nitrospira* LAS18, comammox *Nitrospira* LAS72, and 20 high-quality *Nitrospira* genomes downloaded from the NCBI database. The heatmap displays the average nucleotide identity (ANI) (80–100%) across genomes. Bar plots indicate the GC-content (50–65%) and the total genome length (0–5 Mbp). The analysis focused on core genes shared by all compared *Nitrospira* genomes and singleton genes uniquely possessed by each *Nitrospira* genome.

### Compatible solute synthesis and transporters in canonical *Nitrospira* LAS18, comammox Nitrospira LAS72, and other Nitrospira

The presence of genes for the biosynthesis of compatible solutes (ectoine, hydroxyectoine, proline, trehalose, *L*-D-glutamate, and glycine betaine) and their related transporters, which are involved in adaptation to saline environments, was investigated in 22 *Nitrospira* species, including canonical *Nitrospira* LAS18, comammox *Nitrospira* LAS72, and the 20 high-quality *Nitrospira* genomes retrieved from the NCBI database (**Table 3**). Of note, only comammox *Nitrospira* LAS72 among these 22 *Nitrospira* harbored the *gsmt-sdmt* genes associated with the biosynthesis of glycine betaine. Genes related to ectoine and hydroxyectoine production were not found in any of these 22 *Nitrospira.* All the *Nitrospira* except for LK70 had the genomic potential to produce proline and *L*-D-glutamate. Eleven *Nitrospira*, including LAS18 and LAS72, harbored the functional genes to produce trehalose. Transporter genes for compatible solutes, including glycine betaine, proline, *L*-D-glutamate, and trehalose, for which genetic biosynthesis potentials were identified, were confirmed. Ten *Nitrospira* species, including LAS18 and LAS72, possessed genes related to glycine betaine transport. Among the 10 species, canonical *Nitrospira* LAS18 harbored *opuA* and *opuC*, comammox *Nitrospira* LAS72 contained *opuC* and *opuD*, and *Nitrospira* LK70 had *opuC*, associated with glycine betaine transporters, while the other *Nitrospira* possessed only *proP*, a proline transport gene including glycine betaine (**Table S6**). For trehalose transport, the *msmX* gene encoding multiple sugar-binding transporter ATP-binding protein was found in 7 *Nitrospira* species, including LAS18 and LAS72. None of the 22 *Nitrospira* contained genes relevant for *L*-D-glutamate transport.

**Table 3.**
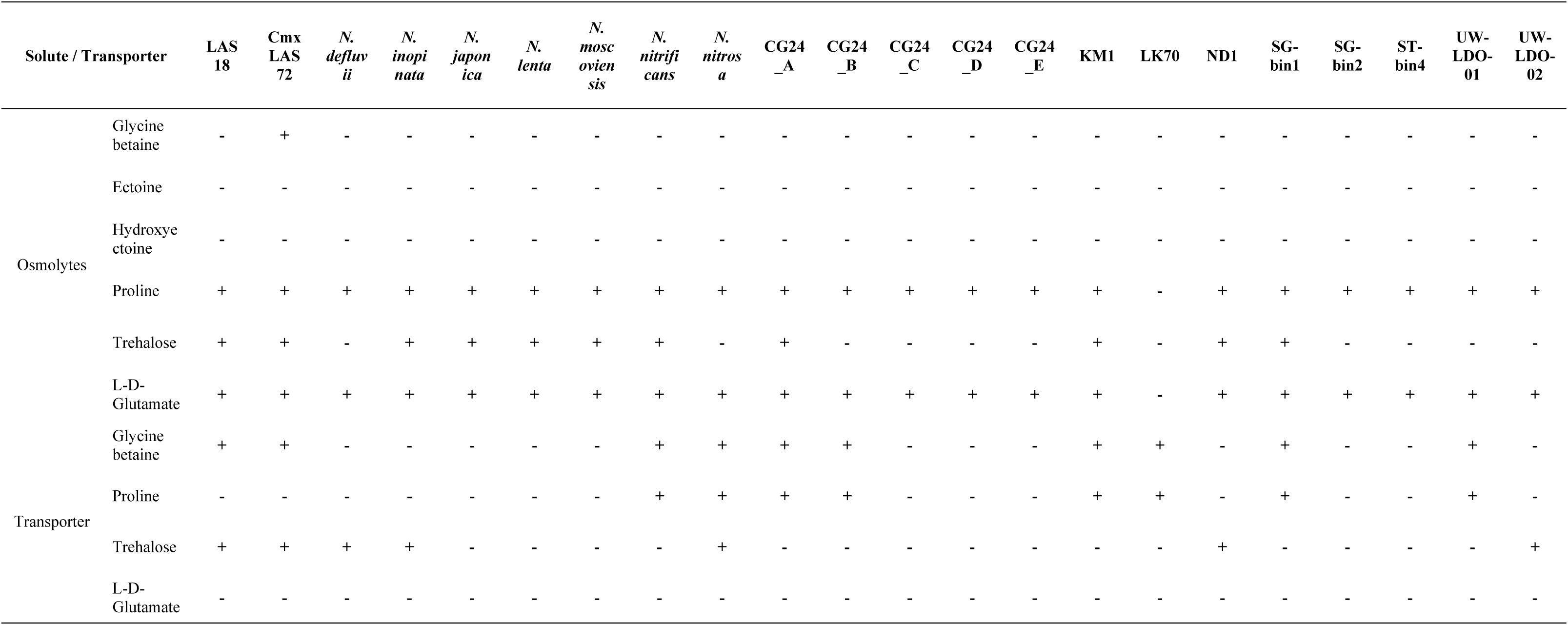
Presence or absence of compatible solutes and transporters for 22 *Nitrospira* genomes including comammox *Nitrospira* LAS72 and canonical *Nitrospira* LAS18.

## Discussion

### Survival strategy and competition between AOM and NOB

In this study, nitrifiers treating ammonia in leachate, *i.e.*, brackish water with high concentrations of ammonia and methanol as an external carbon source, were characterized by a genome-centric approach with hybrid sequencing and integrated binning algorithms. The relative abundance of AOA (1.95%) was similar to or lower than that of the nitrite oxidizer canonical *Nitrospira* LAS18 (2.17%). Dominance of AOA over NOB was reported in a WWTP [63], estuary [64], and soil [65], and for NOB over AOA in marine cycling trickling biofilters [66], culture enriched under oligotrophic soil conditions [67], and a nitrifying SBR operated at low dissolved oxygen concentrations [68]. If comammox *Nitrospira* LAS72 (relative abundance of 0.25%, shown in **Fig. 2**) were included in the AOM in this study, the abundance of AOM (2.20%) would be comparable to that of NOB (2.17%). Although no genes associated with known nitrite transporters were detected in the LAS72 genome (**Tables 2** and **S4**), considering previous studies reporting a transient nitrite accumulation in the bulk liquid of comammox culture systems [69, 70], comammox *Nitrospira* LAS72 might have released nitrite through an unknown mechanism. Subsequently, this nitrite could have been taken up by canonical *Nitrospira* LAS18. On the other hand, a higher abundance of NOB (53.8%) over AOM (7.59%) was reported in a marine recirculating trickling biofilter treating relatively high-concentration NH_4_^+^ (78 mg-N/L), although aerobic nitrite oxidation produces 4.8 times less Gibbs free energy than aerobic ammonia oxidation [66]. The slight difference in the CO_2_ assimilation efficiencies of ammonia and nitrite oxidation (0.077 *vs.* 0.053–0.054 μmol-^14^CO_2_/μmol-NO_2_^−^, respectively), coupled with higher free-energy efficiency of the nitrite oxidation (31%–32% compared to 13% for ammonia oxidation), contributed to NOB dominance over AOM. The higher free energy efficiency likely favored the growth of canonical *Nitrospira* LAS18 in this study, resulting in its slightly higher relative abundance than AOA. Moreover, *Nitrospira* may possess a function to utilize nitrogen source other than nitrite. Koch *et al.* reported the presence of a urease gene in *N. moscoviensis* [71], and canonical *Nitrospira* LAS18 also carries a group of urease and urea transporter genes (**Tables 2** and **S4**). Although urea was not quantified in this study, considering that urea is released by bacterial decay and detritus degradation, converting into ammonia by hydrolysis [71–75], it could potentially be generated in the sedimentation tank after nitrification/denitrification, acting as a potential nitrogen source for canonical *Nitrospira* LAS18. Furthermore, a function possessed solely by canonical *Nitrospira* LAS18 among *Nitrospira* may be a reason for its dominance in this environment. Canonical *Nitrospira* LAS18 was the only *Nitrospira* genome containing the *CLCN2* gene, which encodes a chloride ion channel. The presence of this gene possibly helps LAS18 to cope with a high osmotic pressure triggered by salinity changes, by enabling the absorption and release of chloride ions. In addition, possession of the *GSK3A* gene, pertaining to glycogen synthesis, possibly provides an advantage for survival in starvation conditions after nitrification/denitrification is complete (**Table S5**) [76].

### The presence of comammox *Nitrospira* in this unique environment

To our knowledge, for the first time, a high-quality comammox *Nitrospira* draft genome from a brackish water environment containing a high ammonia concentration was obtained. Comammox *Nitrospira* and AOA compete for ammonia in this environment, and, eventually, AOA (1.95%) prevailed over comammox *Nitrospira* (0.25%). *Candidatus* Nitrosocosmicus LAS21 accounted for 1.62% of the total microbial abundance as the dominant AOA. This species appears more adaptable to higher ammonia concentrations than other AOA [77]. The relative abundance of AOM in this study was comparable to that in several previous studies reporting that comammox *Nitrospira* had a lower relative abundance than that of AOA in full-scale WWTPs [14, 18, 78, 79]. On the other hand, there are controversial reports that comammox *Nitrospira* were dominant in wastewater treatment systems [18, 68, 80–82]. A unique characteristic of the environment in this study stemmed from the presence of methanol, added as an external electron donor, which was plausibly instrumental in favoring the growth of comammox *Nitrospira* LAS72. Although nitrifiers do not possess methanol dehydrogenase, LAS72 is the only *Nitrospira* possessing *fdhA*, which encodes glutathione-independent formaldehyde dehydrogenase, necessary for the metabolism of formaldehyde, a metabolite of methanol. Methanol metabolism was putatively driven in this microbiome by the most dominant methanol-assimilating bacterium, *Methyloceanibacter*, and the formaldehyde produced might be used by comammox *Nitrospira* LAS72. The availability of formaldehyde, which is located upstream of carbon metabolism, may have contributed the preferential growth of comammox *Nitrospira* LAS72. Although the ammonia concentration in the leachate was relatively high (about 130 mg-N/L [22]), the biomass was exposed to nitrogen-limited conditions when ammonia was consumed after nitrification/denitrification during the biomass recycling from the sedimentation tank to the original SBR. Similar to canonical *Nitrospira* LAS18, comammox *Nitrospira* LAS72 harbors the genes for urea transporters and urease; LAS72 is likely to use urea produced by bacterial decay and detritus degradation.

An essential factor for survival in high-salinity conditions is resistance to osmotic pressure [83]. The microbiome in the SBR in this study is required to withstand fluctuations in osmotic pressure caused by the mixing of groundwater and leachate before nitrification/denitrification, and the dewatering after the treatment. Compatible solutes acting as osmoprotectants are essential for microorganisms to remain active in high-salinity environments, and therefore, they synthesize multiple osmolytes in saline environments [84]. Comammox *Nitrospira* LAS72 harbors the genes responsible for production of proline, trehalose, *L*-D-glutamate, and their related transporters. Among the four nitrifiers recovered in this study and other *Nitrospira* previously reported, comammox *Nitrospira* LAS72 is noteworthy because only LAS72 possesses the *gsmt-sdmt* genes, involved in the production of glycine betaine from glycine. Possession of this gene suggests that *Nitrospira* LAS72 is conceivably more adaptable to salinity-fluctuating environments than other comammox *Nitrospira* species.

Our genome-centric analysis illuminates the hidden potential of comammox *Nitrospira* in high-saline and -ammonia conditions, expanding our understanding of the functions of comammox *Nitrospira*. Analysis of gene expression by metatranscriptomics will further improve our understanding of the specific metabolic pathways in comammox *Nitrospira* LAS72 and other nitrifiers, and the cross-feeding relationships between species in this unique methanol-fed environment containing high ammonia and salinity. Furthermore, knowledge gained from future study of brackish environments with different salt and ammonia concentrations, including estuaries and salt lakes, will provide insight into the habitats of comammox *Nitrospira* and their contribution to nitrification.

## Conclusions

This study deciphered the genomes of four nitrifying microorganisms—comammox *Nitrospira* LAS72, canonical *Nitrospira* LAS18, *Candidatus* Nitrosocosmicus LAS21, and *Nitrosarchaeum* LAS73—in an SBR treating high-ammonia brackish landfill leachate with methanol as an added carbon source. The high-quality draft genome for the novel comammox *Nitrospira* LAS72, classified into comammox clade A in lineage II, reveals possible adaptation to this unique environment. Furthermore, canonical *Nitrospira* LAS18 forms a sub-lineage within lineage VII, and is the only representative with a sequenced genome in that sub-lineage. We deciphered the unique metabolic potential of comammox *Nitrospira* LAS72, which alone among *Nitrospira* harbors the *gsmt-sdmt* genes related to the production of glycine betaine from glycine, the *opuD* gene encoding a glycine betaine transporter, and the *fdhA* gene encoding a glutathione-independent formaldehyde dehydrogenase required for formaldehyde metabolism. These unique features possessed by comammox *Nitrospira* LAS72 are likely key factors for it to thrive in brackish and high-ammonia environments. Our discovery of the novel comammox *Nitrospira* LAS72 and the novel sub-lineage VII *Nitrospira* LAS18 broadens our understanding of nitrification mechanisms in saline and high ammonia conditions.

## Competing Interests

The authors declare no competing financial interests.

## Data availability

The sequencing data generated by metagenomic analysis have been deposited in the DDBJ/EMBL/GenBank with the following accession numbers: DRR333436–DRR333437, DRA013194, BioProject: PRJDB12729, and BioSample: SAMD00430153. MAGs for *Candidatus* Nitrosocosmicus LAS21 (LAS21_mss: BQWZ01000001–BQWZ01000031), *Nitrosarchaeum* LAS73 (LAS73_mss: BQXB01000001–BQXB01000005), canonical *Nitrospira* LAS18 (LAS18_mss: BQWY01000001–BQWY01000011), and comammox *Nitrospira* LAS72 (LAS72_mss: BQXA01000001–BQXA01000044) were also deposited.

## Acknowledgements

We thank the late Ms. Kanako Mori for experimental support, Mr. Kuninori Morimoto for bioinformatics analytical support, and Mr. Kousho Hosoi for fruitful discussions. This work was supported by Grants-in-Aid for Scientific Research (20H04362, 22K14348, and 23H03565). Some of this work has been performed using the Center for Computational Science and Engineering of Southern University of Science and Technology. We thank Edanz (https://jp.edanz.com/ac) for editing a draft of this manuscript.

## Notes

### Competing Interest Statement

The authors have declared no competing interest.

