## Supplementary for "Potential survival strategies of novel comammox and nitrite-oxidizing *Nitrospira* present in a reactor treating high-ammonia brackish landfill leachate"

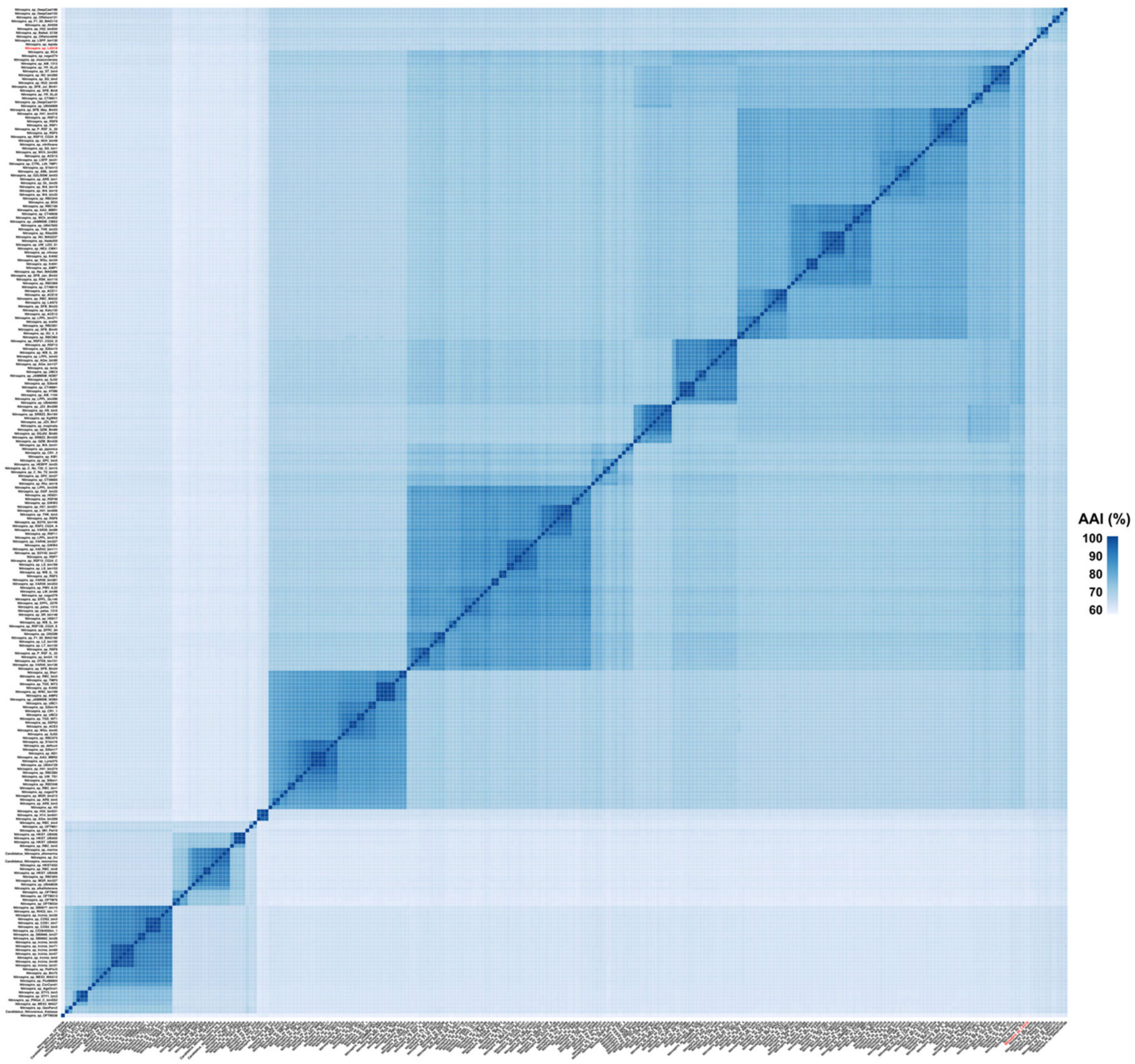


**Fig. S1** Average amino acid identities (AAI) among selected Nitrospira genomes. *Nitrospira sp.* LAS18 is highlighted in red. The heatmap color ranges from dark blue (representing 100% AAI) to white (denoting 55% AAI). *Nitrospira sp.* LAS18 exhibits a range AAI with the rest of *Nitrospira* genomes, from a minimum AAI of 56.7% to a maximum of 69.5%, the latter being with *Nitrospira tepida*.

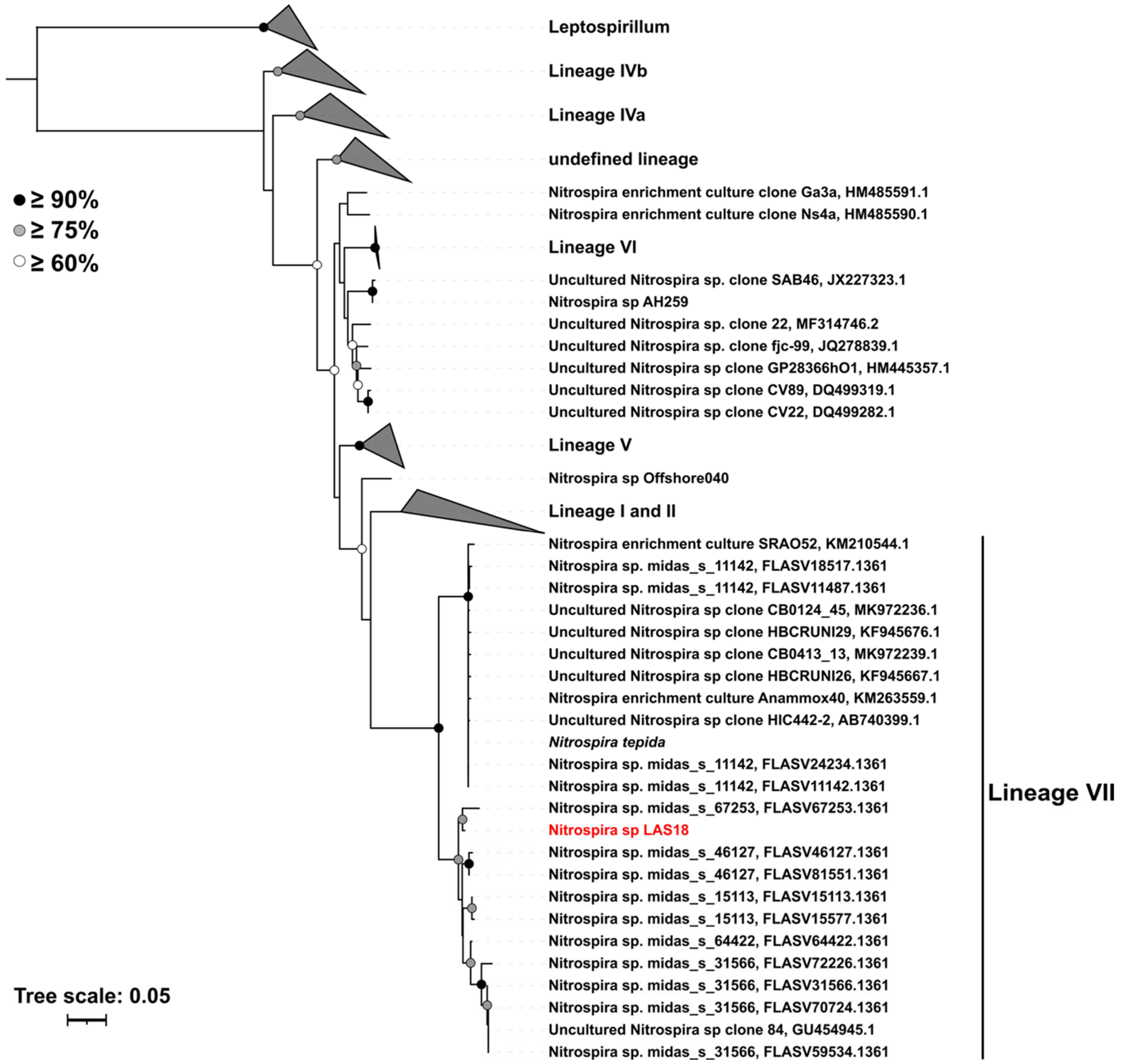


**Fig. S2** Phylogenetic affiliation of *Nitrospira sp.* LAS18 based on the 16S rRNA gene. The maximum-likelihood tree was calculated using MEGAX and 100 bootstrap iterations. All bootstrap support values above 60% are indicated. The analysis involved 168 rRNA gene sequences.

| | CG24_A | CG24_B | CG24_C | CG24_D | CG24_E | KM1 | **LAS_18** | **LAS_72** | LK70 | ND1 | SG_bin1 | SG_bin2 | ST_bin4 | UW_LDO_01 | UW_LDO_02 | defluvii | inopinata | japonica | lenta | moscoviensis | nitrificans | nitrosa |
|---|---|---|---|---|---|---|---|---|---|---|---|---|---|---|---|---|---|---|---|---|---|---|
| CG24_A | 1.00 | 0.85 | 0.85 | 0.84 | 0.84 | 0.85 | 0.87 | 0.86 | 0.84 | 0.84 | 0.83 | 0.84 | 0.84 | 0.86 | 0.84 | 0.84 | 0.85 | 0.83 | 0.84 | 0.83 | 0.84 | 0.84 |
| CG24_B | | 1.00 | 0.85 | 0.84 | 0.83 | 0.84 | 0.91 | 0.83 | 0.84 | 0.84 | 0.84 | 0.84 | 0.85 | 0.83 | 0.83 | 0.82 | 0.85 | 0.83 | 0.84 | 0.82 | 0.88 | 0.83 |
| CG24_C | | | 1.00 | 0.84 | 0.84 | 0.83 | 0.83 | 0.82 | 0.83 | 0.84 | 0.83 | 0.83 | 0.83 | 0.85 | 0.84 | 0.83 | 0.83 | 0.82 | 0.83 | 0.82 | 0.84 | 0.85 |
| CG24_D | | | | 1.00 | 0.85 | 0.83 | 0.86 | 0.83 | 0.83 | 0.83 | 0.83 | 0.83 | 0.83 | 0.83 | 0.83 | 0.84 | 0.82 | 0.83 | 0.89 | 0.84 | 0.84 | 0.83 |
| CG24_E | | | | | 1.00 | 0.82 | 0.86 | 0.82 | 0.84 | 0.84 | 0.83 | 0.84 | 0.83 | 0.83 | 0.83 | 0.83 | 0.83 | 0.83 | 0.83 | 0.83 | 0.82 | 0.83 |
| KM1 | | | | | | 1.00 | 0.87 | 0.89 | 0.85 | 0.85 | 0.83 | 0.82 | 0.83 | 0.87 | 0.85 | 0.85 | 0.86 | 0.85 | 0.84 | 0.83 | 0.86 | 0.86 |
| **LAS_18** | | | | | | | 1.00 | 0.91 | 0.88 | 0.84 | 0.87 | 0.82 | 0.83 | 0.89 | 0.85 | 0.85 | 0.87 | 0.84 | 0.86 | 0.85 | 0.88 | 0.88 |
| **LAS_72** | | | | | | | | 1.00 | 0.83 | 0.86 | 0.83 | 0.84 | 0.83 | 0.84 | 0.87 | 0.86 | 0.87 | 0.86 | 0.86 | 0.84 | 0.83 | 0.84 |
| LK70 | | | | | | | | | 1.00 | 0.84 | 0.84 | 0.83 | 0.83 | 0.83 | 0.85 | 0.86 | 0.85 | 0.83 | 0.83 | 0.84 | 0.84 | 0.83 |
| ND1 | | | | | | | | | | 1.00 | 0.85 | 0.83 | 0.83 | 0.85 | 0.99 | 0.92 | 0.84 | 0.84 | 0.89 | 0.83 | 0.86 | 0.86 |
| SG_bin1 | | | | | | | | | | | 1.00 | 0.84 | 0.84 | 0.83 | 0.83 | 0.83 | 0.84 | 0.83 | 0.83 | 0.83 | 0.84 | 0.83 |
| SG_bin2 | | | | | | | | | | | | 1.00 | 0.95 | 0.83 | 0.83 | 0.83 | 0.83 | 0.83 | 0.83 | 0.83 | 0.84 | 0.83 |
| ST_bin4 | | | | | | | | | | | | | 1.00 | 0.82 | 0.83 | 0.83 | 0.83 | 0.82 | 0.84 | 0.83 | 0.84 | 0.82 |
| UW_LDO_01 | | | | | | | | | | | | | | 1.00 | 0.84 | 0.84 | 0.85 | 0.84 | 0.84 | 0.83 | 0.83 | 0.95 |
| UW_LDO_02 | | | | | | | | | | | | | | | 1.00 | 0.92 | 0.85 | 0.84 | 0.84 | 0.83 | 0.86 | 0.86 |
| defluvii | | | | | | | | | | | | | | | | 1.00 | 0.86 | 0.85 | 0.84 | 0.83 | 0.85 | 0.86 |
| inopinata | | | | | | | | | | | | | | | | | 1.00 | 0.84 | 0.82 | 0.83 | 0.84 | 0.86 |
| japonica | | | | | | | | | | | | | | | | | | 1.00 | 0.83 | 0.83 | 0.84 | 0.87 |
| lenta | | | | | | | | | | | | | | | | | | | 1.00 | 0.83 | 0.84 | 0.85 |
| moscoviensis | | | | | | | | | | | | | | | | | | | | 1.00 | 0.83 | 0.84 |
| nitrificans | | | | | | | | | | | | | | | | | | | | | 1.00 | 0.84 |
| nitrosa | | | | | | | | | | | | | | | | | | | | | | 1.00 |

**Fig. S3** Average Nucleotide Identity (ANI) for 22 *Nitrospira* genomes including comammox *Nitrospira* LAS72 and canonical *Nitrospira* LAS18. ANI with 90% to less than 95% was colored in green, and one with more than 95% was highlighted in orange.

**Table S1** Completeness and contamination for *Nitrospira* genomes. *Nitrospira* genomes downloaded from NCBI meeting with the criteria; completeness > 90% and contamination < 10% were used for comaprison with canonical *Nitrospira* LAS18 and comammox *Nitrospira* LAS72.

| *Nitrospira* species | Completeness [%] | Contamination [%] |
|---|---|---|
| LAS_18 | 95.85 | 3.69 |
| LAS_72 | 93.18 | 3.38 |
| defluvii | 97.67 | 2.27 |
| inopinata | 96.82 | 4.77 |
| japonica | 96.82 | 3.92 |
| lenta | 95.85 | 3.18 |
| moscoviensis | 95.91 | 6.55 |
| nitrificans | 96.76 | 2.73 |
| nitrosa | 96.76 | 2.27 |
| CG24_A | 94.03 | 2.73 |
| CG24_B | 92.22 | 2.78 |
| CG24_C | 92.09 | 2.73 |
| CG24_D | 93.07 | 2.73 |
| CG24_E | 93.98 | 4.09 |
| KM1 | 95.91 | 2.73 |
| LK70 | 93.58 | 5.05 |
| ND1 | 97.67 | 2.73 |
| SG-bin1 | 95.85 | 3.69 |
| SG-bin2 | 95.85 | 3.69 |
| ST-bin4 | 92.06 | 4.6 |
| UW-LDO-01 | 95.8 | 3.64 |
| UW-LDO-02 | 94.36 | 4.66 |

**Table S2** Sequencing data of short and long reads sequenced with DNBSEQ-G400FAST (Short reads) and MinION (Long reads). Pair-read short-read sequencing was implemented twice with two indices to ensure barcode diversity. MinION data shows raw reads with adapters removed and trimmed with Q6.

| Technology | Machine | Read Number | Base Number | Quality cut off [%] | File | Read Length |
|---|---|---|---|---|---|---|
| MGI | DNBSEQ-G400FAST | 19,536,656 | 2,930,498,400 | 89.15 (Q30) | SY1-1_Read1.fq.gz | 150 |
| | | 19,536,656 | 2,930,498,400 | 81.61 (Q30) | SY1-1_Read2.fq.gz | 150 |
| | | 17,582,888 | 2,637,433,200 | 86.53 (Q30) | SY1-2_Read1.fq.gz | 150 |
| | | 17,582,888 | 2,637,433,200 | 80.56 (Q30) | SY1-2_Read2.fq.gz | 150 |
| | | 14,185,213 | 2,127,781,950 | 90.88 (Q30) | SY1-3_Read1.fq.gz | 150 |
| | | 14,185,213 | 2,127,781,950 | 84.41 (Q30) | SY1-3_Read2.fq.gz | 150 |
| | | 12,955,592 | 1,943,338,800 | 89.23 (Q30) | SY1-4_Read1.fq.gz | 150 |
| | | 12,955,592 | 1,943,338,800 | 83.70 (Q30) | SY1-4_Read2.fq.gz | 150 |
| Nanopore | MinION | 3,236,234 | 25,621,131,386 | 81.50 (Q10) | long_read.fastq | 10,593 (N50) |

**Table S3** Hybrid and short read assembly results. All statistics are based on contigs of size >= 500 bp, and "Number of contigs (>= 0 bp)" and "Total length (>= 0 bp)" include all contigs.

| | Hybrid assembly | short read assembly |
|---|---|---|
| Number of contigs (>= 0 bp) | 468,765 | 657,173 |
| Number of contigs (>= 1000 bp) | 69,635 | 131,188 |
| Number of contigs (>= 5000 bp) | 18,768 | 12,526 |
| Number of contigs (>= 10000 bp) | 11,124 | 5,556 |
| Number of contigs (>= 25000 bp) | 4,645 | 2,199 |
| Number of contigs (>= 50000 bp) | 2,207 | 963 |
| Total length (>= 0 bp) | 911,955,543 | 757,919,696 |
| Total length (>= 1000 bp) | 722,902,762 | 481,049,906 |
| Total length (>= 5000 bp) | 625,696,377 | 269,181,616 |
| Total length (>= 10000 bp) | 571,450,752 | 221,687,354 |
| Total length (>= 25000 bp) | 471,150,056 | 170,842,435 |
| Total length (>= 50000 bp) | 386,654,513 | 128,435,148 |
| Number of contigs | 196,482 | 384,890 |
| Largest contig | 4,199,816 | 1,301,673 |
| Total length | 807,556,774 | 653,520,927 |
| GC (%) | 61.8 | 61.4 |
| N50 | 43,606 | 2,719 |
| N75 | 6,670 | 957 |
| L50 | 2,574 | 28,655 |
| L75 | 15,310 | 140,484 |
| Number of N's per 100 kbp | 0.4 | - |

**Table S4** Presence or absence of genes regarding metabolisms without $CO_2$ fixation pathway in Table 2 for four nitrifiers obtained in this study.

| Metabolism | Related genes | *Nitrospira* | | AOA | |
|---|---|---|---|---|---|
| | | canonical_LAS18 | comammox_LAS72 | LAS21 | LAS73 |
| Ammonia oxidation | | - | + | + | - |
| | *amoA* | - | + | - | - |
| | *amoB* | - | + | + | - |
| | *amoC* | - | + | - | - |
| Hydroxylamine oxidation | | - | + | - | - |
| | *hao* | - | + | - | - |
| Nitrite oxidation | | + | + | - | - |
| | *nxrA* | + | + | - | - |
| | *nxrB* | + | + | - | - |
| | *narI* | - | - | - | - |
| Ammonium transporter | | + | + | - | - |
| | *amt* | + | - | - | - |
| | *SLC42A* | - | + | - | - |
| Nitrite/Nitrate transporter | | + | - | - | - |
| | *nrtA* | + | - | - | - |
| | *nrtB* | + | - | - | - |
| | *nrtC* | + | - | - | - |
| | *nrtD* | - | - | - | - |
| | *narK* | + | - | - | - |
| | *nirC* | - | - | - | - |
| | *focA* | - | - | - | - |
| | *focB* | - | - | - | - |
| | *fdhC* | + | - | - | - |
| | *nirC* | - | - | - | - |
| | *narT* | - | - | - | - |
| | *narK* | - | - | - | - |
| Methanol oxidation | | - | - | - | - |
| | *mdh1* | - | - | - | - |
| | *mxaJ* | - | - | - | - |
| | *mxaG* | - | - | - | - |
| | *mdh2* | - | - | - | - |
| | *mxaA* | - | - | - | - |
| | *mxaC* | - | - | - | - |
| | *mxaK* | - | - | - | - |
| | *mxaL* | - | - | - | - |
| | *mxaD* | - | - | - | - |
| | *MOX* | - | - | - | - |
| | *xoxF* | - | - | - | - |
| | *mdh* | - | - | - | - |
| Formaldehyde oxidation | | - | + | - | - |
| | *DAS* | - | - | - | - |
| | *fdhA* | - | + | - | - |
| | *fae* | - | - | - | - |
| | *gfa* | - | - | - | - |
| | *mdo* | - | - | - | - |
| | *fae-hps* | - | - | - | - |
| Nitrite reduction | | + | + | + | + |
| | *nirK* | + | + | + | + |
| | *nirS* | - | - | - | - |
| Nitric oxide reduction | | + | + | + | - |
| | *norB* | - | - | - | - |
| | *norC* | - | - | - | - |
| | *norE* | - | - | - | - |
| | *norD* | + | + | - | - |
| | *norF* | - | - | - | - |
| | *norQ* | + | + | + | - |
| Nitrous oxide reduction | | - | - | - | - |
| | *nosZ* | - | - | - | - |
| Urease | | + | + | - | + |
| | *URE* | - | - | - | - |
| | *ureG* | + | + | - | + |
| | *ureF* | + | + | - | + |
| | *ureD* | + | + | - | + |
| | *ureC* | + | + | - | + |
| | *ureB* | + | + | - | + |
| | *ureA* | + | + | - | + |
| | *ureAB* | - | - | - | - |
| | *E6.3.4.6* | - | - | - | - |
| | *DUR1* | - | - | - | - |
| | *atzF* | - | - | - | - |
| Urea transporter | | + | + | - | - |
| | *urtA* | + | - | - | - |
| | *urtB* | + | - | - | - |
| | *urtC* | + | + | - | - |
| | *urtD* | + | + | - | - |
| | *urtE* | + | + | - | - |

**Table S5** KEGG-annotated singleton genes for (a) canonical *Nitrospira* LAS18 and (b) comammox *Nitrospira* LAS72.

(a)

| K No. | Gene | Name |
|---|---|---|
| K00021 | HMGCR | hydroxymethylglutaryl-CoA reductase (NADPH) [EC:1.1.1.34] |
| K00301 | E1.5.3.1 | sarcosine oxidase [EC:1.5.3.1] |
| K00864 | *glpK* | glycerol kinase [EC:2.7.1.30] |
| K01007 | *pps* | pyruvate, water dikinase [EC:2.7.9.2] |
| K01082 | *cysQ* | 3'(2'), 5'-bisphosphate nucleotidase [EC:3.1.3.7] |
| K01426 | *amiE* | amidase [EC:3.5.1.4] |
| K01535 | PMA1 | H+-transporting ATPase [EC:7.1.2.1] |
| K02635 | *petB* | cytochrome b6 |
| K03215 | rumA | 23S rRNA (uracil1939-C5)-methyltransferase [EC:2.1.1.190] |
| K03258 | *EIF4B* | translation initiation factor 4B |
| K03365 | *FCY1* | cytosine/creatinine deaminase [EC:3.5.4.1 3.5.4.21] |
| K03918 | lat | L-lysine 6-transaminase [EC:2.6.1.36] |
| K05011 | *CLCN2* | chloride channel 2 |
| K06877 | *K06877* | DEAD/DEAH box helicase domain-containing protein |
| K07046 | *K07046* | L-fuconolactonase [EC:3.1.1.-] |
| K07267 | *oprB* | porin |
| K07659 | *ompR* | two-component system, OmpR family, phosphate regulon response regulator OmpR |
| K07925 | RAB39B | Ras-related protein Rab-39B |
| K08331 | ATG13 | autophagy-related protein 13 |
| K08822 | GSK3A | glycogen synthase kinase 3 alpha [EC:2.7.11.26] |
| K08992 | lapA | lipopolysaccharide assembly protein A |
| K09136 | ycaO | ribosomal protein S12 methylthiotransferase accessory factor |
| K09370 | *ISL1* | insulin gene enhancer protein ISL-1 |
| K09373 | LHX2_9 | LIM homeobox protein 2/9 |
| K09378 | *ZFHX3* | zinc finger homeobox protein 3 |
| K10947 | *padR* | PadR family transcriptional regulator, regulatory protein PadR |
| K11648 | *SMARCB1* | SWI/SNF-related matrix-associated actin-dependent regulator of chromatin subfamily B member 1 |
| K12960 | *mtaD* | 5-methylthioadenosine/S-adenosylhomocysteine deaminase [EC:3.5.4.31 3.5.4.28] |
| K16684 | *NF2* | merlin |
| K16841 | *hpxA* | allantoin racemase [EC:5.1.99.3] |
| K17925 | *SNX13* | sorting nexin-13 |
| K19745 | *acuI* | acrylyl-CoA reductase (NADPH) [EC:1.3.1.-] |
| K20331 | *toxA* | toxoflavin synthase [EC:2.1.1.349] |
| K20810 | *mqnX* | aminodeoxyfutalosine deaminase [EC:3.5.4.40] |
| K21310 | *mddA* | methanethiol S-methyltransferase [EC:2.1.1.334] |
| K21677 | *hpnE* | hydroxysqualene dehydroxylase [EC:1.17.8.1] |
| K22083 | *mgsC* | methylamine---glutamate N-methyltransferase subunit C [EC:2.1.1.21] |
| K22522 | LOG | cytokinin riboside 5'-monophosphate phosphoribohydrolase [EC:3.2.2.-] |
| K22747 | *AIFM3* | apoptosis-inducing factor 3 |
| K24273 | *ZRSR* | U2 small nuclear ribonucleoprotein auxiliary factor 35 kDa subunit-related protein |
| K24375 | ZNF687 | zinc finger protein 687 |

(b)

| K No. | Gene | Name |
|---|---|---|
| K00005 | *gldA* | glycerol dehydrogenase [EC:1.1.1.6] |
| K00148 | *fdhA* | glutathione-independent formaldehyde dehydrogenase [EC:1.2.1.46] |
| K05020 | *opuD* | glycine betaine transporter |
| K05643 | *ABCA3* | ATP-binding cassette, subfamily A (ABC1), member 3 |
| K07762 | *PAPPA* | pappalysin-1 [EC:3.4.24.79] |
| K08762 | *DBI* | diazepam-binding inhibitor (GABA receptor modulator, acyl-CoA-binding protein) |
| K10402 | *KIF20* | kinesin family member 20 |
| K10488 | *ZBTB1* | zinc finger and BTB domain-containing protein 1 |
| K10751 | *CHAF1B* | chromatin assembly factor 1 subunit B |
| K11860 | *OTUD7A_B* | OTU domain-containing protein 7 [EC:3.4.19.12] |
| K12472 | *EPS15* | epidermal growth factor receptor substrate 15 |
| K13019 | *wbpI* | UDP-GlcNAc3NAcA epimerase [EC:5.1.3.23] |
| K13866 | *SLC7A4* | solute carrier family 7 (cationic amino acid transporter), member 4 |
| K16148 | *gldM* | alpha-maltose-1-phosphate synthase [EC:2.4.1.342] |
| K16639 | *LACTB2* | endoribonuclease LACTB2 [EC:3.1.27.-] |
| K16923 | *qrtT* | energy-coupling factor transport system substrate-specific component |
| K17294 | *TSPAN4* | tetraspanin-4 |
| K24071 | *gsmt-sdmt* | glycine/sarcosine/dimethylglycine N-methyltransferase [EC:2.1.1.156 2.1.1.157] |
